# Noncanonical targeting contributes significantly to miRNA-mediated regulation

**DOI:** 10.1101/2020.07.07.191023

**Authors:** Jennifer Y. Tan, Baroj Abdulkarim, Ana C. Marques

**Affiliations:** Department of Computational Biology, University of Lausanne, Switzerland

## Abstract

Determining which genes are targeted by miRNAs is crucial to elucidate their contributions to diverse biological processes in health and disease. Most miRNA target prediction tools rely on the identification of complementary regions between transcripts and miRNAs. Whereas important for target recognition, the presence of complementary sites is not sufficient to identify transcripts targeted by miRNAs.

Here, we describe an unbiased statistical genomics approach that explores genetically driven changes in gene expression between human individuals. Using this approach, we identified transcripts that respond to physiological changes in miRNA levels. We found that a much smaller fraction of mRNAs expressed in lymphoblastoid cell lines (LCLs) than what is predicted by other tools is targeted by miRNAs. We estimate that each miRNA has a relatively small number of targets. The transcripts we predict to be miRNA targets are enriched in AGO-binding and previously validated miRNAs target interactions, supporting the reliability of our predictions. Consistent with previous analysis, these targets are also enriched among dosage sensitive and highly controlled genes.

Almost a third of genes we predict to be miRNA targets lack sequence complementarity to the miRNA seed region (noncanonical targets). These noncanonical targets have higher complementary with the miRNA 3’ end. The impact of miRNAs on the levels of their canonical or noncanonical targets is identical supporting the relevance of this poorly explored mechanism of targeting.

## INTRODUCTION

Post-transcriptional regulation by microRNAs (miRNAs) is widespread in eukaryotes (Bartel 2004). These small (20-22 nucleotide) noncoding RNAs guide target recognition by the miRNA-Induced Silencing Complex (miRISC), which in turn leads to transcript degradation or translational repression (Bartel 2004). In animals, loss of function mutations in miRISC proteins (Kataoka et al. 2001; Alisch et al. 2007; Morita et al. 2007; Vasquez-Rifo et al. 2012) or proteins involved in miRNA maturation (Bernstein et al. 2003; Wienholds et al. 2003; Giraldez et al. 2005; Wang et al. 2008) are embryonically lethal. This highlights the importance of post-transcriptional regulation by miRNAs, particularly during early development. In contrast, the phenotypes associated with loss of function mutations in individual miRNAs are often cell/tissue specific, consistent with the spatial and temporally restricted expression of most miRNAs (reviewed in (Bartel 2018)). The penetrance and impact of miRNA loss of function mutations is also highly heterogenous, ranging for example from postnatal lethality (Heidersbach et al. 2013) to increased susceptibility to genetic or environmental stressors (Stadthagen et al. 2013). This heterogeneity is likely a consequence of the diverse functions of genes whose expression is regulated by miRNAs. Given the importance of miRNAs in diverse biological processes in health and disease (Bartel 2018), significant efforts have gone into determining the mechanisms of miRNA targeting and identifying the repertoire of individual miRNA targets (Eulalio and Mano 2015).

In animals, miRNA target recognition often relies on the base pairing between the miRNA seed region (nucleotides 2-8) and target miRNA recognition elements (MREs), frequently found at the target 3’ untranslated regions (UTRs) (Bartel 2004). We refer to this miRNA targeting architecture as “canonical”.

Over the past decades, a wide array of prediction methods were developed to establish the miRNA targetome (Peterson et al. 2014). These softwares primarily rely on identifying well-established canonical signatures of miRNA:target interactions, including sequence complementarity, accessibility, location and conservation of MREs, as well as favourable binding energy between miRNA and target (Peterson et al. 2014). Notably, the overlap between predictions made by different softwares is often small (Sethupathy et al. 2006) and fewer than 1% of predicted miRNA targets are supported by experimental evidence for miRISC binding (Fridrich et al. 2019). These analyses highlight the extent of false predictions and our poor understanding of the properties that underlie efficient targeting and regulation by miRNAs. Machine learning approaches, in particular those guided by well-defined interactions between miRNA and target (McGeary et al. 2019), have the potential to improve computational prediction of miRNA targets (Schafer and Ciaudo 2020). But given the scarcity of large and comprehensive sets of true miRNA targets, the development of such tools is currently restricted.

Alternatively, miRNAs targets can be experimentally identified based on the evidence of their interaction with miRISC or that their expression levels change in response to genetic miRNA perturbations. Given the ease to measure the impact of miRNA level changes on gene expression levels transcriptome-wide, the latter approach has been extensively used (Bartel 2018). However, the impact of miRNA perturbation on gene expression is 1) often smaller than the inter-individual variation in transcript levels, suggesting many of these changes are inconsequential (Pinzon et al. 2017); and 2) this approach does not allow the distinction between changes due to miRNA targeting (direct) or miRNA-driven changes in the levels of gene expression regulators, including transcription factors or RNA binding proteins (indirect) (Thomas et al. 2010).

Cross-liking Immunoprecipitation (CLIP) based approaches have been used to identify transcripts bound by miRISC (reviewed in (Lin and Miles 2019)) *in vitro* (i.e. (Hafner et al. 2010)) and *in vivo* (i.e. (Li et al. 2019)). Experimental evidence for AGO-binding greatly increases the accuracy of miRNA target predictions. However not all transcripts bound by miRISC are miRNA targets (Agarwal et al. 2015). Furthermore, whereas, as expected, transcripts with reproducible binding are enriched among true targets, they are depleted in lowly abundant transcripts leading to false negative predictions (Wessels et al. 2019).

Despite their low sensitivity, methods like CLASH (Helwak et al. 2013) or miR-CLIP (Imig et al. 2015) have experimentally detected direct miRNA:target interactions and revealed that a sizable fraction of these interactions occurs at regions outside of the 3’UTRs or rely on noncanonical or mismatch containing seed sites (Helwak et al. 2013). High-throughput analysis of affinity binding also supports that some miRNAs can recognize their targets through noncanonical configurations (Becker et al. 2019; McGeary et al. 2019). The extent of noncanonical targeting cannot be assessed using most of the currently available computational prediction tools, further highlighting their limitations.

To overcome some of limitations of the current miRNA target prediction methods, we developed an approach that explores genetically driven changes in gene expression between human individuals. This approach allowed us to identify canonical and noncanonical targets that respond to physiological changes in miRNA levels.

## RESULTS

### Identification of miRNA targets in LCLs

To identify physiologically relevant miRNA-target interactions, we leveraged small RNA sequencing for a panel of 360 lymphoblastoid cell lines (LCLs), derived from healthy individuals of European descent with known genotype (Lappalainen et al. 2013; 1000 Genomes Project et al. 2015). For 62% (n=444) of all LCL expressed miRNAs, we identified at least one single nucleotide polymorphisms (SNPs) in the vicinity of their corresponding miRNA gene whose changes were significantly (FDR<0.05, Methods) associated with mature miRNA levels (Figure 1A, Supplementary S1A). We refer to these variants as miRNA quantitative trait loci (mirQTLs). We reasoned that targets of these miRNAs should be inversely associated with these mirQTLs *in trans* (Methods), consistent with miRNAs being negative regulators of gene expression. We used polyA-selected RNA-sequencing data in these cells and identified 6325 mRNAs (42.6% of the 14847 LCL-expressed genes) associated in *trans* in the inverse direction with at least one miRNA mirQTL (n=6430 joint eQTLs, Methods, Figure 1B,C, Supplementary S1B).

**Figure 1.**
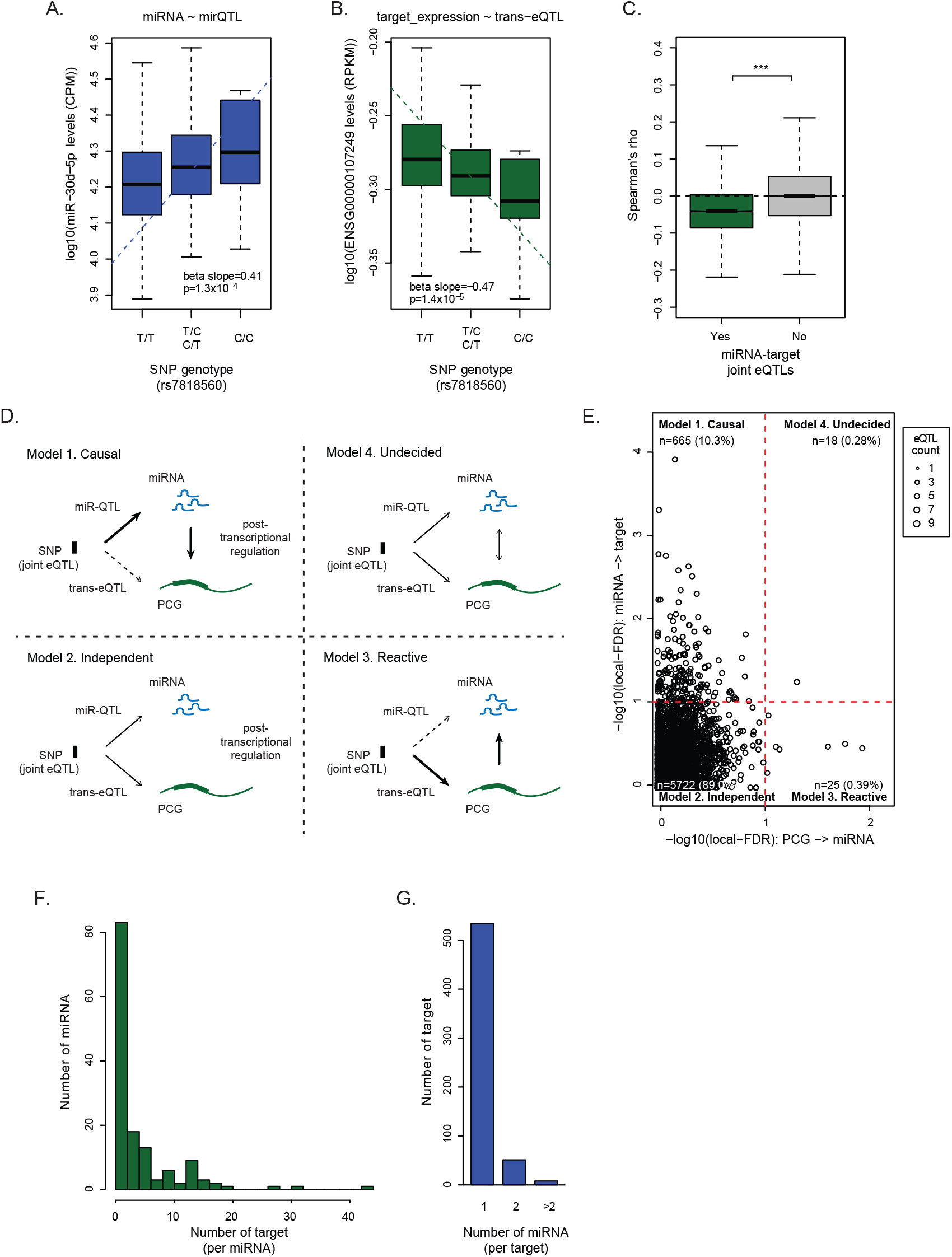
GenVar approach detects physiologically relevant miRNA targets. (A) Distribution of miR-30d-5p expression (Y-axis, CPMs) in individuals with different genotypes of the SNP variant (rs7818560, X-axis, mirQTL). (B) Distribution of GLIS3 (ENSG00000107249) expression (RPKM, Y-axis) in individuals with different genotypes of the SNP variant (rs7818560, X-axis, *trans*-eQTL). Beta slope and pvalue of eQTL associations are shown in the inset. (C) The distribution of co-expression (Spearman’s correlation) between levels of miRNAs and genes that are jointly (green) or not jointly (grey) associated to the same SNP variant. Differences between groups were tested using a two-tailed Mann-Whitney *U* test. *** *p* < 0.001. (D) Schematic representation of the four models of causal inference testing that predict the relationship between joint eQTL variant (black box) associations with the miRNA levels (mirQTL) (blue) and target gene expression level (*trans*-eQTL) (green): (1) direct association between the variant and miRNA levels mediates the indirect association between that and target gene expression (causal model); (1) the variant is independently associated with miRNA and target levels (independent model); (3) direct association between the variant with target expression mediates the indirect association between that and miRNA levels (reactive model); and (4) the interaction between miRNA levels and target expression is complex (undecided model). Direct associations are depicted as solid lines and indirect associations as dash lines. (E) Scatterplot depicting causal inference testing local FDR associated with each the four models (as illustrated in D). The likelihood of target regulating miRNA levels is plotted on the x-axis and the likelihood of miRNA regulation target expression is plotted on the y-axis. Number and proportion of joint eQTLs are provided in brackets for each model. Dotted red lines denote significance threshold at local FDR < 0.1. Frequency distribution of the (F) number of predicted targets per miRNA and (G) number of targeting miRNAs per gene.

The association between a mirQTL and a mRNA can be a consequence of 1) the mRNA levels being regulated by the miRNA (causal model); 2) the miRNA and mRNA being independently associated with the same variant, for example if the same transcription factor regulates transcription of both the miRNA and mRNA (independent model); 3) the miRNA being regulated by the mRNA (reactive model); or 4) an undetermined mode of regulation (undecided model, Figure 1D)(Wang and Michoel 2017). We used causal inference analysis to distinguish between these different possibilities for each joint eQTL-miRNA-mRNA trio. We found that with a false discovery rate of 10%, a tenth (10.3%, n=665) of all associations are mediated by changes in miRNA levels (Figure 1E). In total, we identified around 4% of all LCL-expressed genes (593 out of 14847) to be causal targets for 143 miRNAs. This corresponds to an average of 5 targets per miRNA (median 2 targets, ranging between 1 and 44) (Figure 1F) and 1-2 miRNAs targeting each transcript (Figure 1G, Supplementary Table ST1).

Given stronger mirQTL associations might bias towards detecting cases where miRNA levels causally mediate the association between the variant and target expression, we compared the overall strength of associations between mirQTLs and target *trans*-eQTLs and found no evidence that stronger mirQTLs would favour causal model predictions (Supplementary Figure S1C). Hereafter we refer to transcripts predicted to be targeted by miRNAs based on their genetic variations in humans as GenVar-targets.

### Limitations of GenVar-target identification approach

It is well established that the power to detect quantitative trait loci for relatively lowly expressed genes is limited (GTEx Consortium et al. 2017). We assessed the extent by which this hampered our ability to use the GenVar approach to predict miRNA targets. miRNAs with predicted GenVar-targets (n=143) are among the most highly expressed in LCLs (n=716, p=9.4×10^−7^, two-tailed Mann-Whitney U test, Figure 2A). Similarly, we found GenVar-targets are more highly expressed than non-targets (p=6.57×10^−5^, two-tailed Mann-Whitney U test, Figure 2B). These findings possibly reflect the approach’s limitation to predict lowly expressed miRNAs and target transcripts.

**Figure 2.**
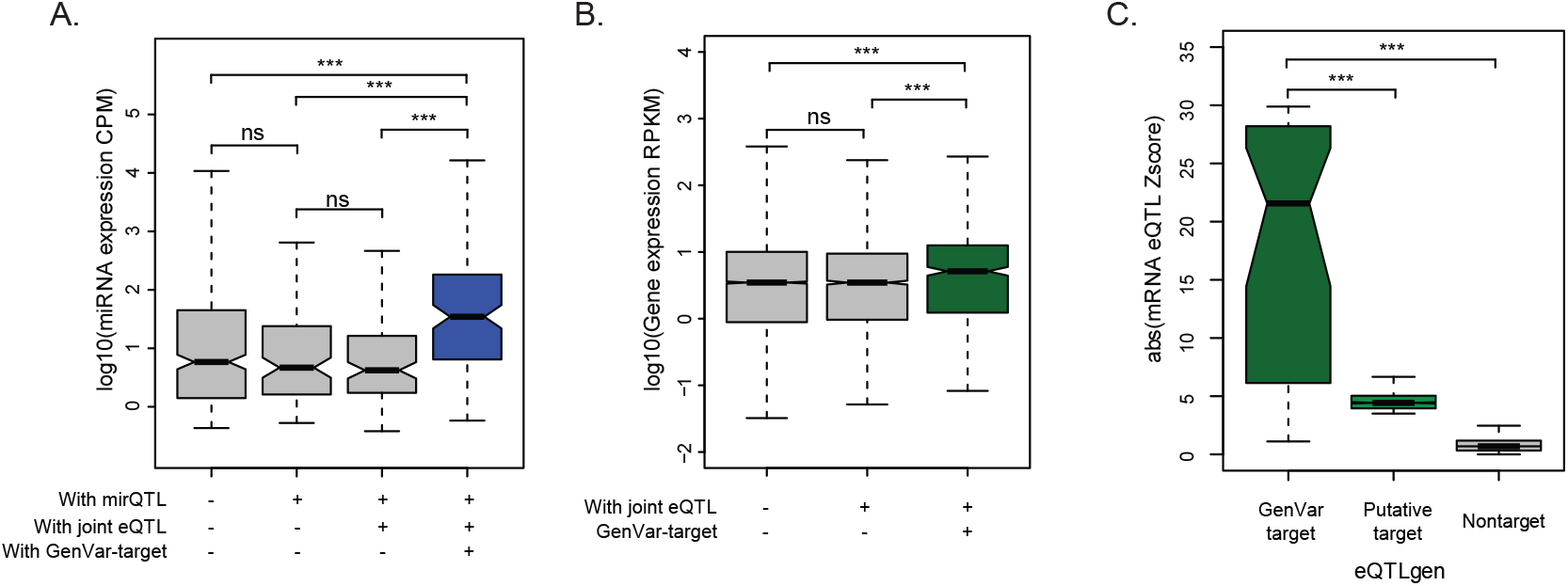
Limitations of the GenVar approach. (A) Distribution of subsets of LCL-expressed miRNA levels with mirQTL associations, whose mirQTLs are jointly associated with target expression (joint-eQTLs), and with predicted GenVar-targets (blue). (B) Distribution of subsets LCL-expressed gene expression levels with those that are jointly associated with a mirQTL (joint-eQTL), and those that are predicted GenVar-targets (green). (C) In blood samples, distribution of *trans*-eQTL associations (Zscore) between validated GenVar-targets (dark green), putative targets (light green) or nontargets (grey) with mirQTLs. Differences between groups were tested using a two-tailed Mann-Whitney *U* test. * *p* < 0.05; ** *p* < 0.01; *** *p* < 0.001; NS *p* > 0.05.

Cohort size is also a well-established contributor to the power of quantitative trait analysis (Beavis 1998). To assess whether our predictions were limited by this, we used a dataset that is almost 100 times larger (31,684 blood samples, eQTLGen consortium (Võsa et al. 2018)) than the one used to identify GenVar-targets (360 LCLs (Lappalainen et al. 2013)). As the expression levels for mature miRNAs is unavailable for these blood samples and most blood expressed miRNAs are also detected in LCLs (Juzenas et al. 2017), we assumed mirQTLs are conserved between the two cohorts. We used publicly available *trans*-eQTL data from eQTLgen to assess miRNA-target associations in the blood samples. This data is available for a subset of human variants (GWAS-associated SNPs, n=10,317) and includes 101 miRQTL variants associated for 23 miRNAs with at least one GenVar-targets in LCLs (total 137 targets). In eQTLgen blood samples, 66% of GenVar-targets (90/137) were also significantly associated in *trans* and in the same direction with their respective miRNA, supporting the interaction between them. This high replication rate support robustness of the association between miRNAs and GenVar-targets. Consistent with this, replicated *trans*-eQTL associations between GenVar-targets and mirQTLs in blood are significantly stronger compared to non-targets in LCLs (p<2.2×10^−16^, two-tailed Mann-Whitney U test, Supplementary Figure S2A). Compared to replicated associations between mirQTLs and GenVar-targets, those only found in LCLs (34%) were similarly associated in blood (p=0.335, two-tailed Mann-Whitney U test, Supplementary Figure S2A), suggesting some of these are likely explained by LCL-specific miRNA regulation.

In addition to the GenVar-targets we validated in blood (validated GenVar-targets), we found the expression levels of another 48 genes are also correlated in *trans* with mirQTLs for 9 of the 23 miRNAs (median 1 target per miRNA, ranging from 1 to 22). Moreover, we also detected 470 putative targets for 23 miRNAs with no predicted GenVar-targets in LCLs (median 4 targets per miRNA, ranging from 1 to 53). We refer to these 518 genes as putative targets. However, despite their *trans*-association with mirQTLs, since miRNA levels are not available in eQTLgen, we could not infer the causal relationship between miRNAs and these putative targets. Given in LCLs, GenVar-targets only accounted for a tenth of all genes associated with mirQTLs (Causal model, Figure 1E), we predict only a small portion of these putative targets are regulated by miRNAs. This hypothesis is supported by the significantly stronger *trans*-associations, in blood, between mirQTLs and validated GenVar-targets compared to putative targets (p<2.2×10^−16^, two-tailed Mann-Whitney U test, Figure 2C). Nevertheless, our findings suggest that the small LCL cohort size likely limited our power in comprehensively detecting all miRNA targets.

### miRNA GenVar-targets are enriched in AGO-binding and experimentally validated targets

Using three widely-used miRNA prediction softwares: TargetScan (Agarwal et al. 2015), RNA22 (Miranda et al. 2006) and miRWalk (Sticht et al. 2018), we estimated between 88% and 93% of mRNAs to be targeted by at least 1 of the 143 GenVar miRNAs (Supplementary Figure S3A). The proportion of genes (n=593) we predicted to be targeted by these miRNAs in LCLs is at least 20 times smaller (Supplementary Figure S3A). The small number of miRNA targets might not be surprising given the low fraction of miRNA-target predictions common to all the 3 tools (11-21%, n=92812 out of 474,114, 825,295, 448,787 total predictions from TargetScan, RNA22, and miRWalk, respectively, Supplementary Figure S3B) and the previously described high rate of false positives associated with the different prediction methods (Fridrich et al. 2019).

The extent of these differences and our approach’s limitations probed us to investigate how different target prediction sets associate with properties of true miRNA targets. We found that the expression of GenVar-targets is more strongly inversely correlated with the levels of their targeting miRNAs compared to targets predicted by other computational tools in human LCLs (Figure 3A).

**Figure 3.**
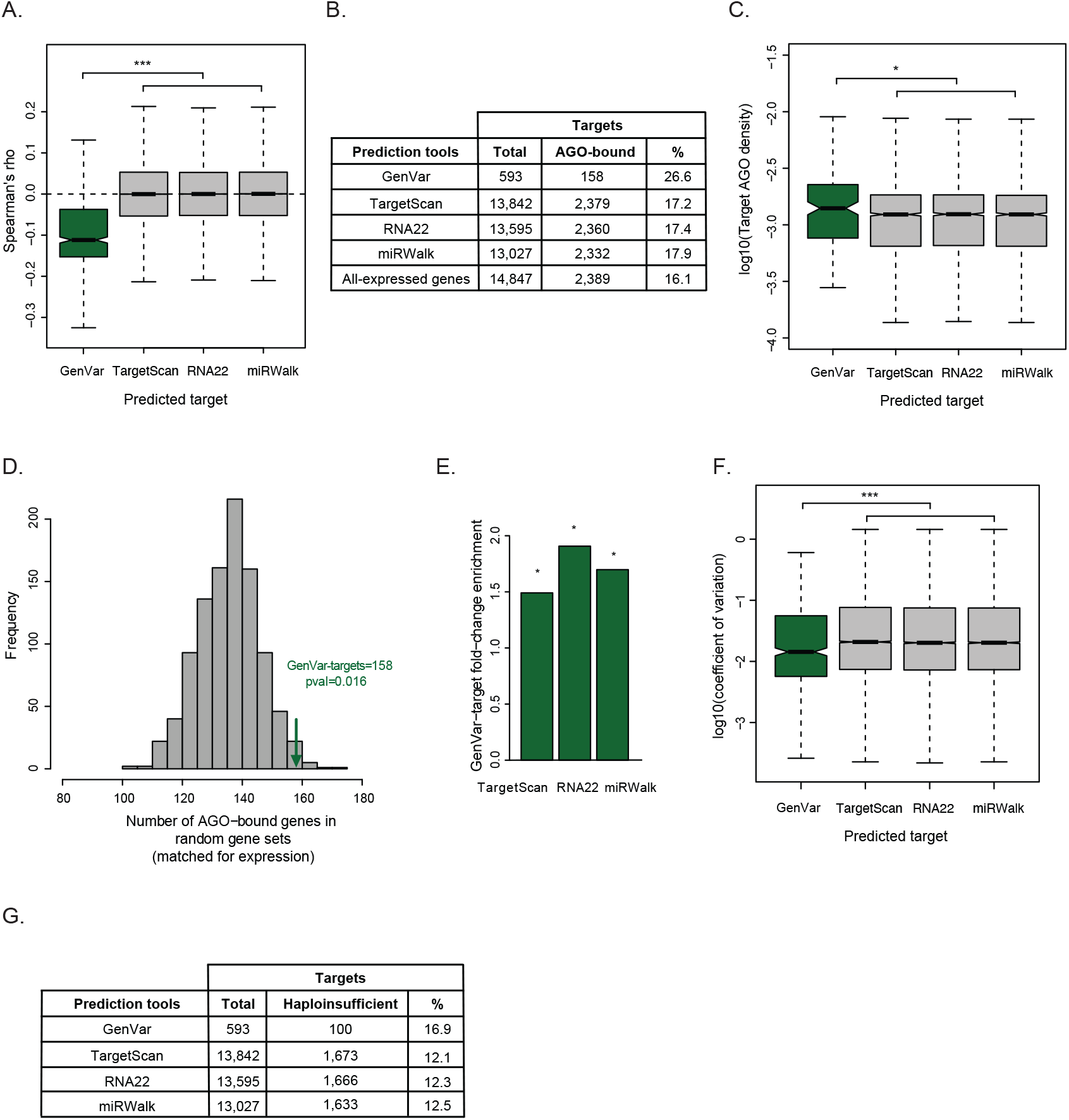
GenVar targets are a reliable set of miRNA targets. (A) Distribution of co-expression (Spearman’s correlation) between miRNA levels and their GenVar-targets (green) or targets predicted by three other tools (TargetScan, RNA22, and miRWalk, grey). (B) Table of the number and proportion of GenVar-targets or targets predicted by other tools to be bound by AGO2. (C) Distribution AGO2 binding density (length of AGO2 binding site/target 3’ UTR length) of GenVar-targets (green) and targets predicted by other tools. (D) Distribution of the number of AGO2-bound genes within 1000 sets of randomly sampled genes from other predicted targets with matching expression levels and 3’ UTR length as GenVar-targets. Green arrow represent the number of AGO2-bound GenVar-targets. (E) Of all predictions by each of the three prediction tools, the enrichment in the proportion of experimentally validated (using direct Luciferase assays) GenVar-target predictions over nonGenVar-target predictions. (F) Distribution of the coefficient of variation (CV) in expression levels of GenVar-targets (green) and targets predicted by other tools (grey). (G) Table of the number and proportion of target predictions that are annotated as haplo-insufficient. Differences between groups were tested using a two-tailed Mann-Whitney *U* or two-tailed Fisher’s exact test. * *p* < 0.05; ** *p* < 0.01; *** *p* < 0.001; NS *p* > 0.05.

Using publicly available iCLIP data in LCLs (Wan et al. 2014), we determined the extent by which the transcripts predicted by the different approaches were bound by AGO2. To minimize the impact of unspecific AGO2 binding in these analyses, we only considered transcripts with experimental evidence of binding across at least two thirds (4 out of 6) of the sequenced LCL samples (Methods). We found that GenVar-targets are significantly more often bound by AGO2 (26.6%) than genes predicted by the other 3 tools (<17.9%, p<2.8×10^−7^, two-tailed Fisher’s exact test, Figure 3B). AGO2-binding density is also significantly higher at GenVar-targets compared to other predicted targets (p<0.02, two-tailed Mann-Whitney U test, Figure 3C), supporting the enrichment of GenVar-targets in miRISC binding and miRNA regulation.

As expected, transcripts with detectable AGO2 binding in iCLIP data are in general more highly expressed (p<2.4×10^−10^, two-tailed Mann-Whitney U test, Supplementary Figure S4A), reflecting the technical limitations of the technology (Wessels et al. 2019). More than 75% (n=447/593) of GenVar-targets are less abundant than the median expression level of transcripts with experimental evidence of AGO2-binding (n=1101/14847, Supplementary Figure S4B), suggesting that AGO2-binding at many GenVar-targets may have escaped detection due to their relatively low expression. To account for this bias, we compared the extent of AGO2 binding at GenVar-targets relative to randomly sampled expression-matched genes predicted by the other 3 tools and we found that GenVar-targets are significantly more often bound by AGO2 than their counterparts from other prediction methods (1.17-1.19 fold enrichment, p<0.024, permutation test, Figure 3D, Supplementary Figure S4C-F).

We tested the proportion of GenVar-targets that have been experimentally validated. Given the known ascertainment bias to experimentally validate previously computationally predicted miRNA targets, we compared the proportion of experimentally validated predictions by each of the three tools which are also detected using the GenVar approach to those not predicted by GenVar. We found a significantly higher fraction of GenVar-targets (p<0.05, two-tailed Fisher’s exact test) has been experimentally validated using direct luciferase assays (1.5-1.9 fold-enrichment, Figure 3E, Supplementary Figure S4G) and in all catalogued experiments (1.4-1.7 fold-enrichment, Supplementary Figure S4H,I) relative to targets predicted by other tools (Karagkouni et al. 2018), further corroborating that our approach enriches for real miRNA targets.

miRNA-mediated repression on most genes is frequently lower than the inter-individual variability in their expression levels (Pinzon et al. 2017). This observation led to the proposal that only the levels of genes whose expression is under tight control, such as haplo-insufficient genes, can be effectively regulated by miRNAs. Consistent with this, we found that the variation in GenVar-target expression across individuals in the human population are significantly lower compared to target predicted made by other tools (p<9.6×10^− 5^, two-tailed, Mann-Whitney U test, Figure 3F). We also found that GenVar-targets are enriched in haplo-insufficient genes (p<3.0×10^−3^, two-tailed Fisher’s exact test, Figure 3G), supporting the notion that miRNA-mediated phenotypical changes are likely due to their ability to regulate a relatively small set of dose-sensitive genes (Pinzon et al. 2017; Seitz 2017).

### Cancer-disrupted miRNA abundance is associated with causal target expression changes

Given the limitations in iCLIP technique and the ascertainment bias in reporting of miRNA validation experiments, we sought an alternative unbiased approach to assess the reliability of GenVar-target predictions. We took advantage of extensive genotyping and transcriptomic data from blood-derived cancer samples to assess the impact of changes in miRNA expression on GenVar-target levels. Specifically, we identified individuals that carry copy number variants (CNVs) that overlap primary miRNA loci in Lymphoid Neoplasm Diffuse Large B-cell Lymphoma (DLBC, Figure 4A)(Cancer Genome Atlas Research et al. 2013). These CNVs resulted in homozygous loss or multiple amplifications of 35 miRNAs (48 samples in total, median 1 samples per CNV, Supplementary Table ST2).

**Figure 4.**
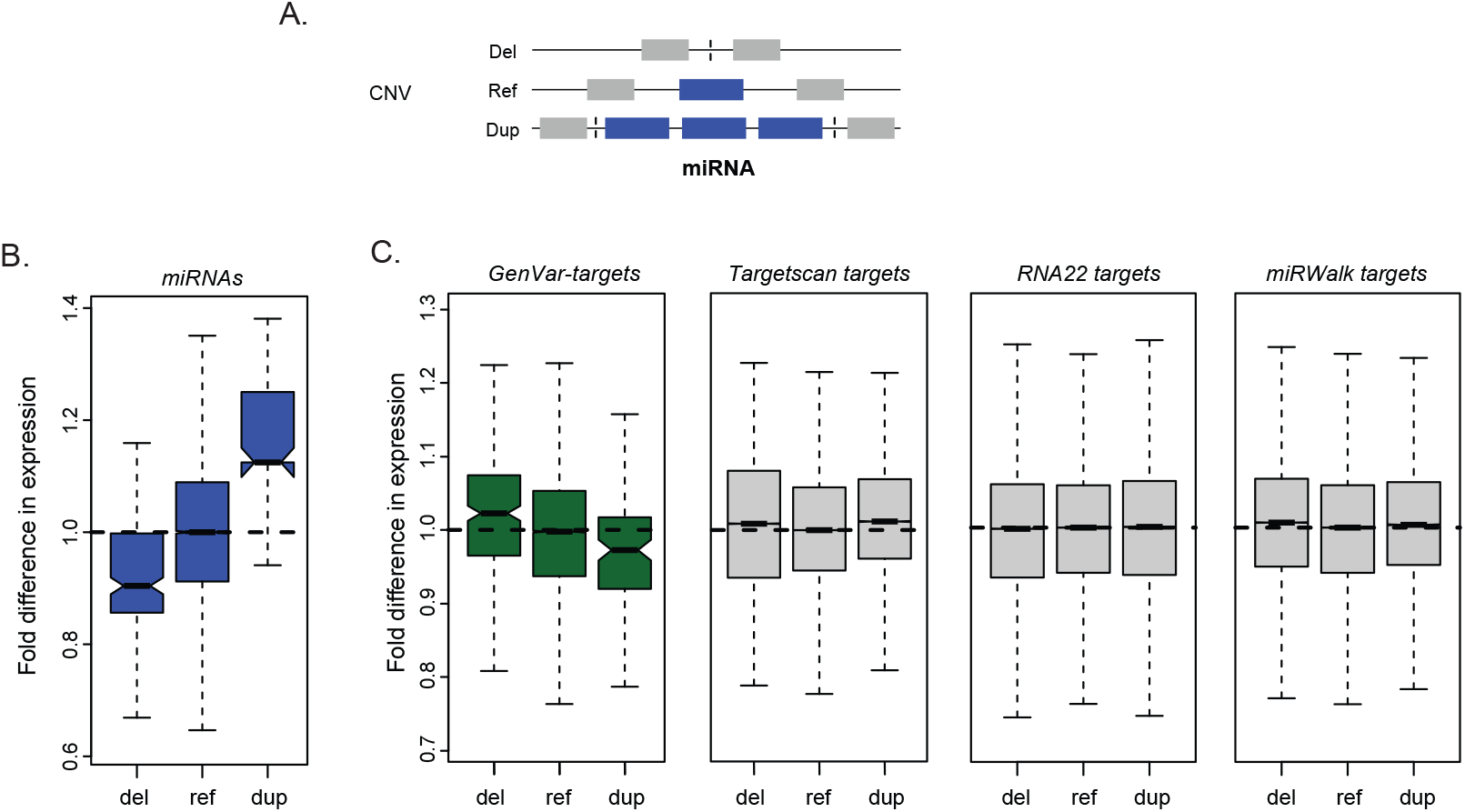
GenVar-target levels are impacted by change in miRNA copy number. (A) Copy number variations in DBLC cancer samples can result in duplications or deletions of primary miRNA loci. (B) Distribution of the fold difference in miRNA levels (CPM, blue), relative to the median of diploid samples, of individuals that carry homozygous deletions, diploid genotype, or multiple duplications at primary miRNA loci. (C) Distribution of fold difference in gene expression levels (RPKM) of GenVar-targets (green) or targets predicted by other tools (grey), relative to the median of diploid samples, between individuals that carry homozygous deletions, diploid genotype, or multiple duplications at their targeting miRNAs. Differences between groups were tested using a two-tailed Mann-Whitney *U* test. * *p* < 0.05; ** *p* < 0.01; *** *p* < 0.001; NS *p* > 0.05.

Mature miRNA expression levels corroborated that, as expected, these CNVs led to decreased or increased miRNA expression by at least 9 and 12%, respectively (Figure 4B) relative to individuals that are diploid for the miRNA gene. The relative small impact of copy number changes on miRNA expression are likely a consequence of samples containing mixed normal and tumor cells (Cancer Genome Atlas Research et al. 2013). Changes in miRNA abundance were significantly negatively associated with expression levels of predicted GenVar-targets (p<0.05, two-tailed Mann-Whitney U test) but not with the levels of targets predicted by other tools (Figure 4B,C), substantiating direct miRNA post-transcriptional regulation of GenVar-targets.

In summary, our results support that GenVar-targets are enriched 1) for AGO2-binding, 2) for experimentally validated miRNA-target interactions, and 3) are inversely impacted by changes in disrupted miRNA levels in cancer samples. These observations support that GenVar-targets represent a reliable catalogue of biologically relevant miRNA targets.

### miRNA targeting is enriched in canonical binding at 3’UTRs

Given that our approach does not rely on the identification of perfectly complementary regions to the miRNA seed within transcripts, we next sought to estimate the extent of canonical and noncanonical miRNA targeting (Hausser and Zavolan 2014). Consistent with previous reports (Lewis et al. 2005), we found the majority of GenVar-targets (404 out of 593, 68.1%) harbour perfect seed matches for their targeting miRNAs at their 3’ UTRs (canonical targets)(Figure 5A). This proportion is significantly higher than randomly sampled genes with matching expression level and 3’ UTR length (p=0.001, permutation test, Supplementary Figure S5A). Compared to other predicted targets that also contain complementary seed regions, alignment profiles of canonical miRNA GenVar-target interactions showed these transcripts bear longer sequence complementary binding sites to miRNA seed region (p=0.039, two-tailed Mann Whitney test, Supplementary Figure S5B,C). This is consistent with significantly higher number of 8-mer and 7-mer binding sites found within GenVar-targets (52.5% 6-mer, 30.2% 7-mer, and 17.3% 8-mer) compared to other predicted targets (p<4.3×10^−3^, two-tailed Fisher’s exact test, Supplementary Figure S5D).

**Figure 5.**
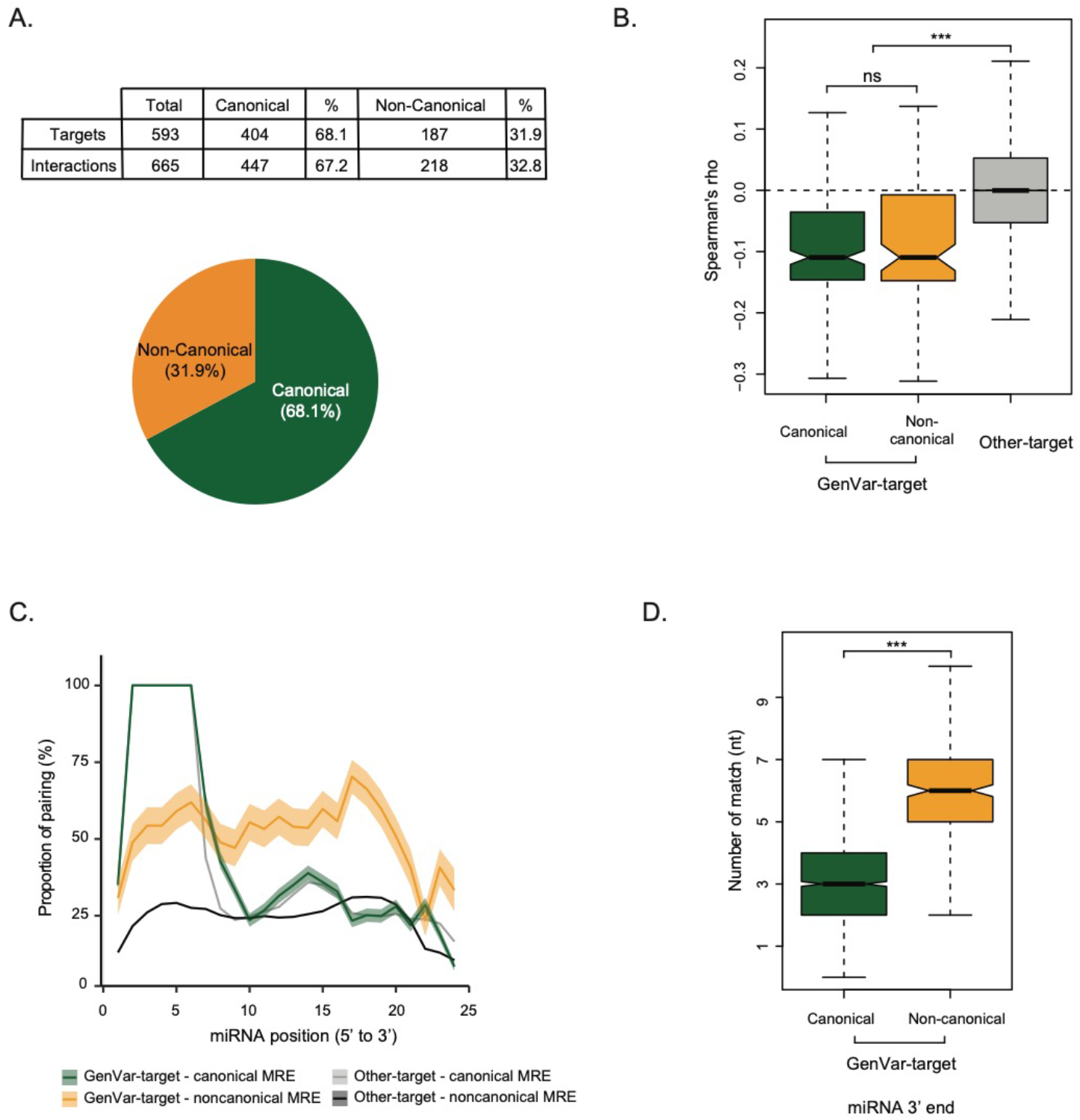
A sizeable fraction of GenVar-targets is noncanonical. (A) Table (Upper panel) and pie chart (lower panel) illustrating the number and proportion of canonical and noncanonical GenVar-targets. (B) Distribution of co-expression (Spearman’s correlation) between miRNA levels and their canonical (green) and noncanonical (yellow) GenVar-targets, as well as targets predicted by other tools (grey). (C) Binding profile illustrating the average fraction of complementary sequence alignment between target and miRNA at each nucleotide position across the mature miRNA (5’ to 3’) for canonical and noncanonical GenVar-targets (green and yellow, respectively), as well as targets predicted other tools that contain canonical and noncanonical binding sites (light and dark grey, respectively). (D) Distribution of the number of complementary sequence alignments (nt) between canonical (green) or noncanonical (yellow) GenVar-targets with 3’ end of miRNAs. Differences between groups were tested using a two-tailed Mann-Whitney *U* test. * *p* < 0.05; ** *p* < 0.01; *** *p* < 0.001; NS *p* > 0.05.

In addition, some miRNAs have also been found to interact with regions in their target’s 5’ UTRs or coding sequences (CDSs) (Chi et al. 2009; Helwak et al. 2013). Of the 31.9% of GenVar-targets (n=189) with no canonical miRNA binding sites in their 3’ UTR targets, seed-matching sites were found at their 5’ UTR and CDS for 9.0% (n=17) and 31.2% (n=59), respectively. Unlike canonical GenVar-targets (Supplementary Figure S5A), these proportions are not significantly different from randomly sampled genes with matching expression level and length of 5’ UTR or CDS (p>0.09, permutation test, Supplementary Figure S5E,F). Our findings are in line with the current understanding that miRNA targeting at 3’ UTR is more likely to lead to physiological changes in target abundance (Gu et al. 2009).

### Noncanonical targets bear extensive complementarity to miRNA 3’ end

Apart from perfect matches to the miRNA seed region that form canonical miRNA binding sites, sequence complementarity sites outside the seed region have also been shown to contribute to guiding miRNA target interactions (Becker et al. 2019; McGeary et al. 2019). To investigate binding architecture that underlie noncanonical miRNA targeting, we focused on the 31.9% of GenVar-targets that lack seed matching miRNA binding sites at 3’ UTRs (Figure 5A, Supplementary Table ST3). In LCLs, the correlation between miRNAs and their canonical or noncanonical targets is statistically indistinguishable (p=0.81, two-tailed Mann-Whitney U test, Figure 5B). Similar impact on canonical and noncanonical GenVar-target expression was also observed as a result of disrupted miRNA levels by CNVs in blood cancer samples (p>0.29, two-tailed Mann-Whitney U test, CNVs disrupted the levels of 34 and 21 miRNA canonical and noncanonical GenVar-targets, respectively, Supplementary Figure S6). These findings support that other features may contribute to determining miRNA-mediated regulation for some miRNAs. Compared to randomly sampled genes with matching expression level and length, noncanonical GenVar-targets were not enriched to contain seed-matching MREs at 5’ UTR or CDS (p>0.05, permutation test, Supplementary Figure S7A,B).

We assessed the alignment between noncanonical GenVar-targets and their associated miRNAs. Using adequate alignments between 68% of target 3’ UTR and their respective miRNAs, we found each miRNA have on average between 2-3 noncanonical targets (ranging from 1 to 20, Supplementary Figure S7C). Of all miRNAs with GenVar-targets, 43.3% regulate gene expression only through the canonical mode of binding, whereas 20.3% have only noncanonical targets, and 36.4% have a mixture of both (Supplementary Figure S7D). Alignment profiles of noncanonical miRNA GenVar-target interactions showed that, as expected, complementarity to miRNA seed regions is significantly less frequent than that found for canonical targets (p<2.2×10^−16^, two-tailed Mann Whitney test, Figure 5C, Supplementary Figure S7E). In contrast, extensive nucleotide complementarity was observed between noncanonical targets and 3’ end of miRNA (p<2.2×10^−16^, two-tailed Mann-Whitney test, Figure 5C,D), suggesting imperfect seed interaction may be compensated by additional binding outside of miRNA seed region.

## DISCUSSION

Target identification remains one of the biggest challenges when trying to understand the contribution of miRNAs to post-transcriptional regulation of gene expression.

Here we propose a target prediction method that leverages genetically driven changes in miRNA and transcript expression levels within humans to identify miRNA targets (GenVar-targets). Compared to targets predicted by other approaches, GenVar-targets are enriched in AGO2 binding, supporting their frequent association with the miRISC. As expected, given their established role in post-transcriptional repression, miRNA copy number changes in cancer, are inversely correlated with GenVar-target expression. This is in contrast to what we observe for miRNA targets predicted using other approaches, whose expression remain unchanged. Finally, a sizable fraction of GenVar-targets are validated in an independent dataset. These observations support the physiological relevance of our miRNA target predictions and the high specificity, especially relative to available alternatives, of the GenVar approach.

GenVar-targets account for 4% of mRNAs expressed in lymphoblastoid cell lines (LCLs). The proportion of transcripts we predict to respond to physiological changes in miRNA levels is significantly smaller than what is estimated by other tools. This discrepancy between ours and other estimates is in part explained by the relatively high rate of false positive predictions of other methods [(Fridrich et al. 2019) and supported by our own analysis]. We also analyzed how different factors, including the expression levels of miRNAs, mRNAs and size of the human cohort, impact the sensitivity of the GenVar approach and likely increase false negative predictions. We found that, as expected, miRNA target associations are significantly stronger in larger cohorts. The large cohort size also allowed us to identify an additional set of putative targets not detected using the smaller set of LCL samples. We therefore anticipate that the ongoing trend to expand population cohort size will result in increased prediction sensitivity and has the potential to extend the repertoire of physiologically relevant miRNA targets.

In summary, our analysis supports the functional relevance of GenVar-targets with relatively low false positive predictions. High specificity of GenVar predictions is accompanied by decreased sensitivity that we anticipate can be minimized with increased cohort size. Furthermore, given most miRNAs are tissue specifically expressed, GenVar-targets identified in LCLs are therefore most likely relevant for this and closely related cells/tissues. The application of this approach to other cells or tissues has the potential to uncover the miRNA targetome in different tissues and to reveal potential tissue-specific features of miRNA targeting.

We further investigated the miRNA target binding architecture of GenVar predictions. Specifically and given that the GenVar approach does not require prior knowledge of miRNAs binding features, it provides an unbiased target set to investigate the extent of noncanonical targeting by miRNAs. Consistent with previous studies (Chi et al. 2009; Hafner et al. 2010; Helwak et al. 2013), we found a sizable fraction (around 32%) of miRNA targets are noncanonical. The impact of miRNA on target expression is independent of binding architecture since variation in miRNA levels was similarly associated with changes in the expression of their canonical or noncanonical targets. Our observations are in line with recent high-throughput binding affinity experiments (Becker et al. 2019; McGeary et al. 2019) and earlier studies (Vella et al. 2004; Didiano and Hobert 2006), and support the functional relevance of noncanonical targeting by miRNAs and the diversity of their binding architecture.

This mode of targeting, which we predict accounts for around a third of miRNA target interactions, is missed by most prediction methods and thus remains poorly understood. Specifically, we found that most noncanonical targets sites that lack perfect seed matches contained additional sequence complementarity with 3’ ends of the miRNA, suggesting mismatches between target and miRNA seed regions can be compensated by additional matches outside the seed. The availability of this high confidence set of transcripts regulated by miRNAs opens avenue to study the role of additional molecular entities in miRNA regulation.

## MATERIAL AND METHODS

### Mapping of molecular quantitative trait loci (QTLs)

We used miRNA abundance and processed genotype for human EBV-transformed lymphoblastoid cell lines (LCLs) derived from 360 healthy individuals of European descent (CEU, GBR, FIN and TSI) (downloaded from EBI ArrayExpress, accession E-GEUV-1)(Lappalainen et al. 2013). We tested the association between single nucleotide polymorphisms (SNPs) located within the 1Mb of primary miRNA transcript loci for each of the 715 miRNAs expressed in LCLs with available expression data (Lappalainen et al. 2013). Only SNPs with minor allele frequency (MAF) greater than 5% were considered in the QTL analyses. miRNA eQTL associations were estimated using FastQTL (version 2.184) (Ongen et al. 2016). To assess the significance of the correlations globally, we first applied multiple testing correction on FastQTL estimated associations by permuting the expression levels of each miRNA 1000 times and noted the maximum permuted absolute regression coefficient (r_max_). We considered only miRNA eQTLs with an observed absolute regression coefficient (r_obs_) greater than 95% of all permuted r_max_ values to be significant (Lappalainen et al. 2013). We further performed Benjamini-Hochberg multiple testing correction to estimate FDR (<5%) for all SNPs within the 1Mb vicinity of pri-miRNAs. We identified significant miRNA-eQTLs (mirQTLs) for 62% (n=444) of LCL-expressed miRNAs.

The expression levels (RPKM) of protein-coding genes (n=14,847) in the same 360 LCL samples were downloaded from a previous study (as described in (Tan et al. 2017)). We identified *trans*-eQTLs between all expressed genes and mirQTLs using the same approach as described above with the exception that multiple-testing correction was applied for all *trans*-eQTL tested. Genes in the vicinity (within 1Mb) of pri-miRNAs were not considered in the analysis. Of the 14,847 protein-coding genes expressed in LCLs, we found 6325 genes (42.6%) to be associated in *trans* and in the inverse direction with mirQTLs for 444 miRNA (6430 joint eQTLs).

### Causality inference between miRNA abundance and target protein-coding gene expression

To infer the causal relationship between miRNA levels and putative target gene expression, we considered joint-eQTLs that are associated with both miRNA levels (mirQTL) and their putative target gene abundance (*trans*-eQTL) (6430 joint eQTLs associated with 444 miRNAs and 6325 protein-coding genes). For all such triplets of joint-eQTL – miRNA – target gene expression, we performed causal inference testing using a Bayesian approach as implemented by Findr (Wang and Michoel 2017) by testing the models: (1) the causal model with miRNA where regulates gene expression; (2) the independent model where joint-eQTL variants are independently associated with miRNA levels and gene expression; (3) the reactive model where gene expression mediates miRNA abundance; and (4) the undecided model where causative relationship between miRNA and gene is more complex (Wang and Michoel 2017). Findr was run using the recommended test (findr.pijs_gassist) with default parameters as described in (Wang and Michoel 2017) and significant associations (FDR<10%) were considered in the analysis.

### Assessing sensitivity and replication rate of the GenVar approach

To assess sensitivity of the GenVar approach, we estimated the number of GenVar-targets undetected using LCL samples (n=360, Geuvadis)(Lappalainen et al. 2013) in a considerably larger cohort of blood samples (n=31,684 (Võsa et al. 2018)). Given mature miRNA levels are not available in the blood samples, for LCL-identified mirQTL with *trans*-eQTL data available in blood (64 miRNA, 101 mirQTLs), we considered *trans*-eQTL associations between all protein-coding genes and these mirQTL to identify additional miRNA targets that are missed in LCLs in blood. We applied multiple testing correction (Benjamini-Hochberg) for the number of *trans*-eQTL variants tested and identified additional miRNA GenVar-targets as those with FDR<5% and are undetected in LCLs.

Robustness of *trans*-eQTL associations identified in LCLs was also assessed in the large cohort of blood samples (Võsa et al. 2018). We considered all *trans*-eQTL associations between GenVar-targets and mirQTLs tested in blood (association data is available in blood for 23 mirQTL variants for 39 miRNAs with GenVar-targets identified in LCLs). For these *trans*-eQTLs, we applied multiple testing correction (Benjamini-Hochberg adjusted p-value <5%) for the number of variants tested. We considered associations found in the same direction in eQTLgen blood samples as that found in Geuvadis LCLs to be replicated.

### Comparison with existing miRNA target prediction tools

We downloaded predictions from TargetScan (version 7) (Agarwal et al. 2015), RNA22 (version 2) (Miranda et al. 2006), miRWalk (version 3) (Sticht et al. 2018) and filtered for miRNAs and protein-coding genes expressed in LCLs.

### Enrichment in AGO2 binding

We assessed the interaction of GenVar-targets with the miRISC complex using publicly available iCLIP data to identify regions bound by AGO (Wan et al. 2014) (GSE50676). AGO-bound transcripts were downloaded from GSE50676. We only considered AGO binding sites consistently detected in at least two thirds of all replicates (4 out of 6 samples). In total, out of all LCL-expressed genes considered in our analysis (n=14,847), 18% (n=2,669) were considered to be robustly bound by AGO.

To account for differences in expression levels, LCL-expressed genes predicted as miRNA targets by other tools were divided into two sets of 4 equally sized bins based on their expression levels. For each GenVar-target, a gene is randomly selected without replacement from their expression-match gene bins.

### Overlap with experimental validated targets

We used a database containing experimentally validated miRNA target interactions (DIANA-TarBase v8) (Karagkouni et al. 2018) to evaluate the proportion of validated GenVar-targets. We filtered for interactions between LCL-expressed miRNAs and target genes and we considered only interactions validated using all validation experiments (n=386,278 interactions) or direct luciferase assays (n=190,334 interactions). To avoid ascertainment bias towards enriched experimental validation of previously computationally predicted targets, when assessing the experimentally validated fraction of GenVar-targets, we considered GenVar-targets that overlapped predictions from other tools and compared these to predictions from other tools that were not detected by GenVar. 463, 171 and 219 GenVar-targets were also predicted by TargetScan, RNA22, and miRWalk, respectively. Most predictions from other tools were not predicted by GenVar as targets (474,069, 825,124 and 448,568 for TargetScan, RNA22, and miRWalk, respectively).

### Overlap with haplo-insufficient genes

Haplo-insufficient genes were downloaded from the DECIPHER database (v10)(Firth et al. 2009) and from (Dang et al. 2008). We merged annotations from the two studies. In total, 1754 LCL-expressed genes were found to be haplo-insufficient (1577 from DECIPHER and 288 from Dang et al. 2008).

### Impact of genetic variation at miRNA loci on GenVar-target expression

We considered all copy number variations (CNVs) that overlap primary miRNA transcript loci from blood-derived cancer samples (Lymphoid Neoplasm Diffuse Large B-cell Lymphoma, DLBC) with available genotyping and transcriptomic data (Cancer Genome Atlas Research et al. 2013). We identified CNVs within 48 individuals that led to homozygous loss or multiple amplifications of 35 miRNAs predicted to have at least one GenVar-target (each deep gain or loss event is found in a median of 1/48 samples). CNV data was downloaded from the TCGA Pan-Cancer cohort on the Xena Functional Genomics Explorer (Goldman et al. 2019). For miRNAs disrupted by CNV, we plotted the average distribution of the fold difference in miRNA levels and their respective target gene expression for each DLBC sample, relative to the median expression of individuals that are diploid for the miRNA gene.

### Identification of noncanonical miRNA targets

We considered all GenVar-targets with seed-matching (including all 8mer, 7mer and 6mer sites) miRNA recognition elements (MREs) at their 3’ UTR, as predicted by the standalone version of TargetScan (version 7) (Agarwal et al. 2015), as canonical targets. Comparisons of seed-matching MREs were made to other canonical MRE-containing targets predicted by TargetScan. Seed-matching MREs located at transcript 5’ UTR and CDS were predicted using the standalone version of TargetScan (version 7) (Agarwal et al. 2015). Ensembl annotated transcript sequences were used in the analysis (version 98). We tested the enrichment of MRE frequency found at 5’ UTR/CDS/3’ UTR against that found at 1000 sets of randomly sampled LCL-expressed genes with matching expression level and respective transcript segment length. LCL-expressed genes were divided into two sets of 4 equally sized bins based on their expression levels or length (5’ UTR, CDS, or 3’ UTR), independently. For each canonical GenVar-target, a gene is randomly selected without replacement from the intersection of their expression-match and length-match gene bins.

For noncanonical GenVar-targets, we predicted miRNA binding sites by aligning the full sequence of their targeting mature miRNA with the 3’ UTR of target genes using RNAduplex using default parameters with added parameters allowing all alignments within 5kcal/mol of the optimal structure to be considered for downstream filtering and GU pairs not considered in the analysis (RNAduplex -e 5 --noGU) (ViennaRNA Package 2.0 (Lorenz et al. 2011)). Alignments that contained consecutive gaps in miRNA sequence or multiple gap openings in target sequence were discarded given the miRISC is unlikely to bind to targets that would result in consecutive bulges in binding (Brennecke et al. 2005). No restriction was set on the number of mismatches in alignment.

### Statistical tests

All statistical analyses were performed using the R software environment for statistical computing and graphics (R Development Core Team 2008). When multiple comparisons were made, we reported the highest p-value of all comparisons.

## Data Access

Analyses were performed using publicly available command-line tools. Versions and deviations from parameters used are as detailed in the Methods. All scripts used to parse the results are available upon request.

## Acknowledgements

We thank members of the Marques group for valuable comments and discussion. We thank Zoltán Kutalik and Olivier Delanau for discussion on population genomics analysis. We thank Constance Ciaudo for discussion on miRNA target validation. This work is funded by the Swiss National Science Foundation grant (PP00P3_179065 to ACM).

## Authors’ contributions

JYT, BA and ACM designed the study. JYT and BA performed the experiments and analyzed the results. JYT, BA and ACM discussed the results. ACM supervised the study. JYT and ACM wrote the manuscript. All authors approved the manuscript.

