## Supplementary figures and images for "Noncanonical targeting contributes significantly to miRNA-mediated regulation"

### Supplementary Figure S1

Supp Figure S1.

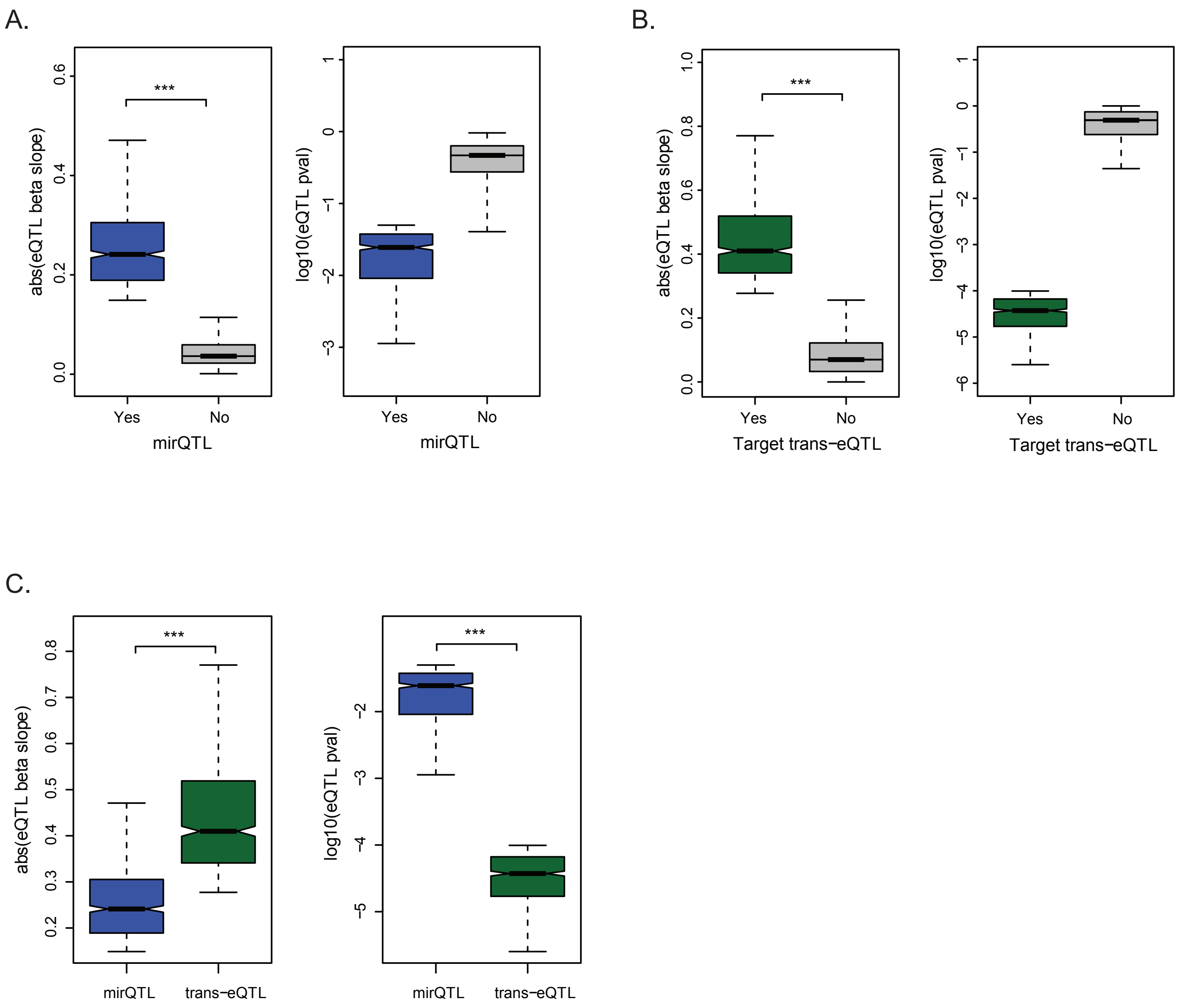

### Supplementary Figure S2

Supp Figure S2.

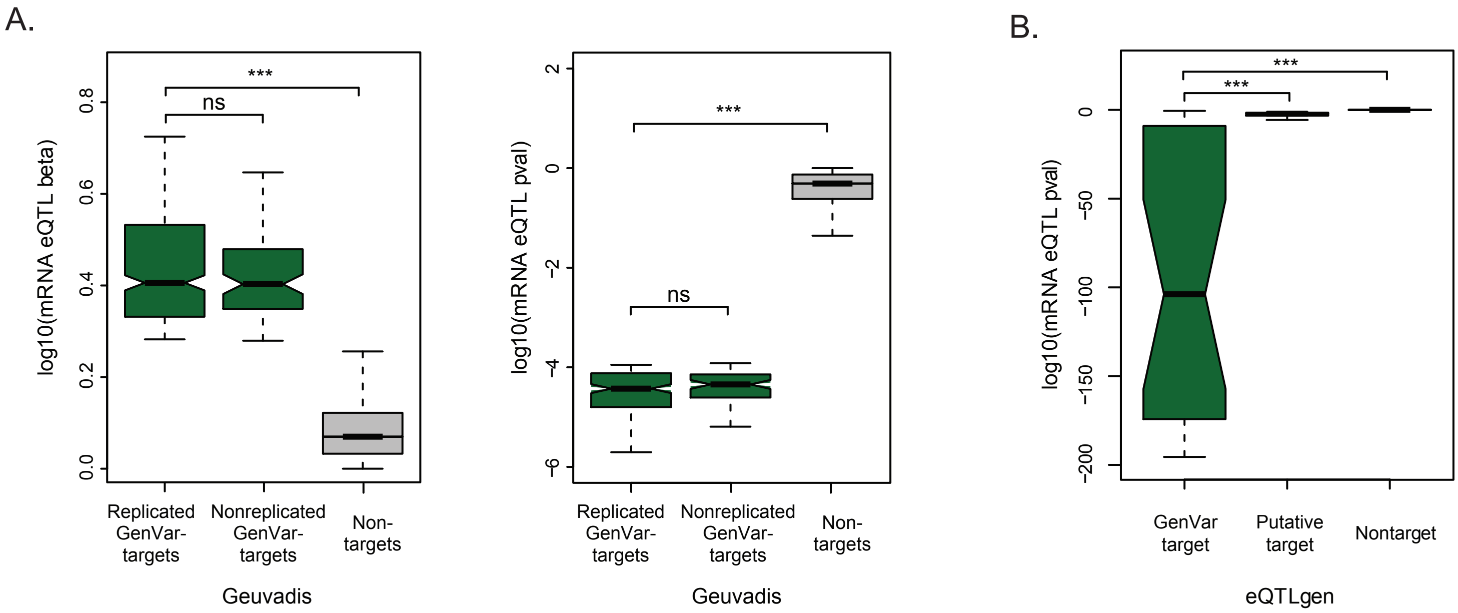

### Supplementary Figure S4

Supp Figure S4.

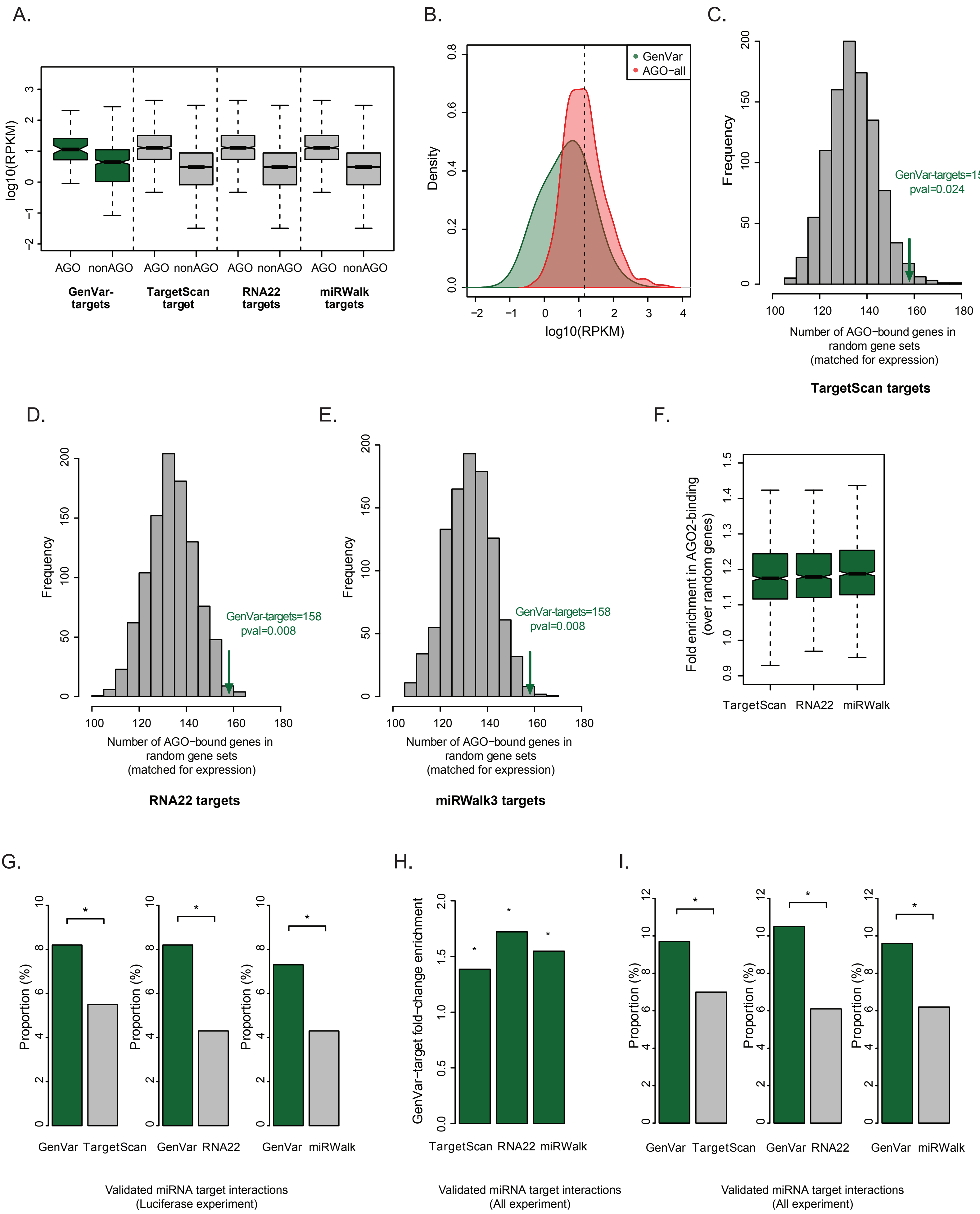

### Supplementary Figure S5

Supp Figure S5.

A. B. C.

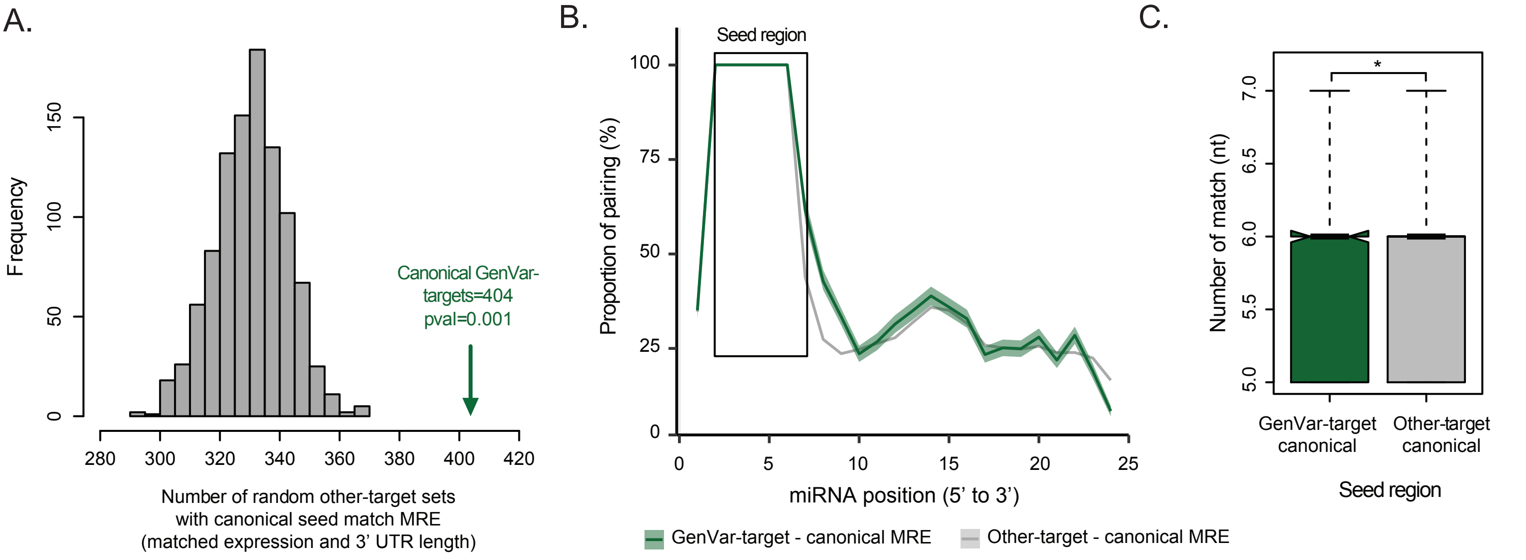


D.

| | Canonical binding interactions | | | |
|---|---|---|---|---|
| | Genvar-target | % | Other-target | % |
| 8mer | 109 | 17.3 | 192142 | 13.6 |
| 7mer | 190 | 30.2 | 345709 | 24.4 |
| 6mer | 331 | 52.5 | 879102 | 62.0 |

E. F.

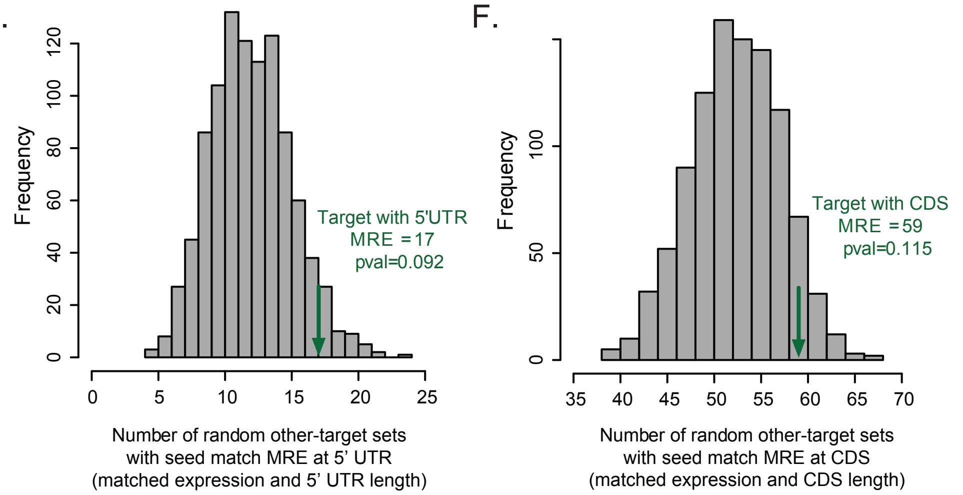

### Supplementary Figure S6

Supp Figure S6.

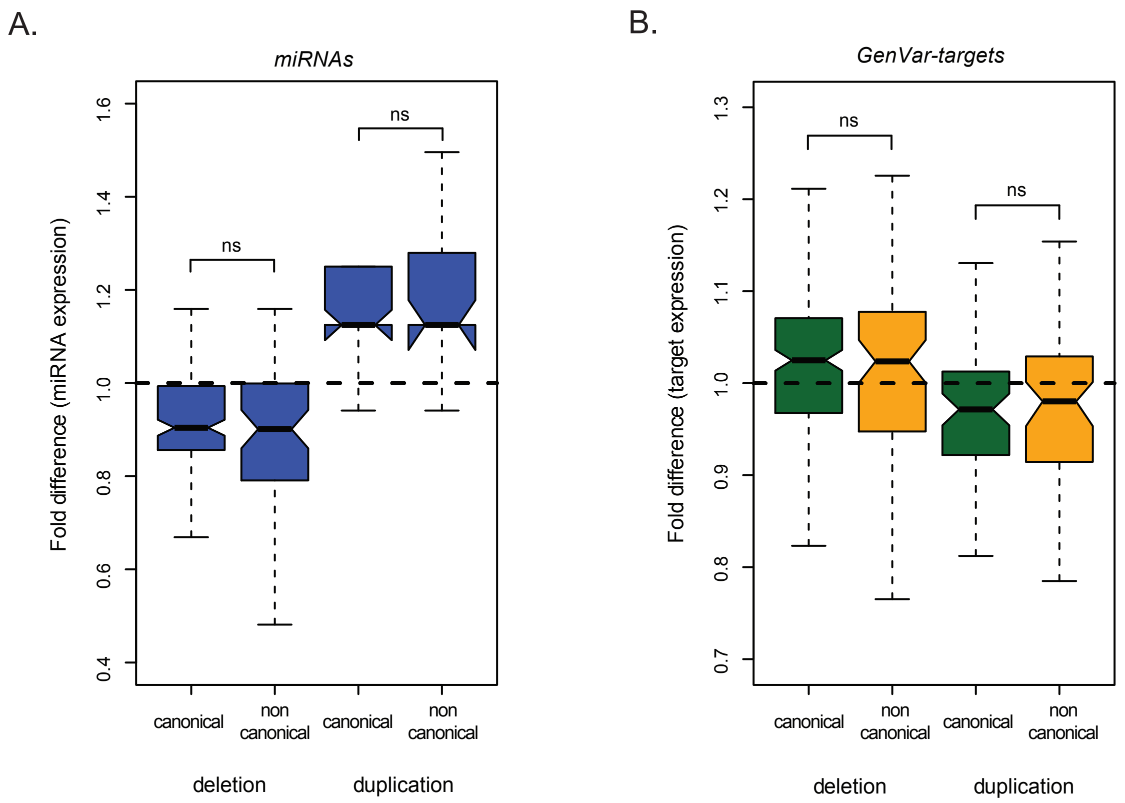

### Supplementary Figure S7

Supp Figure S7.

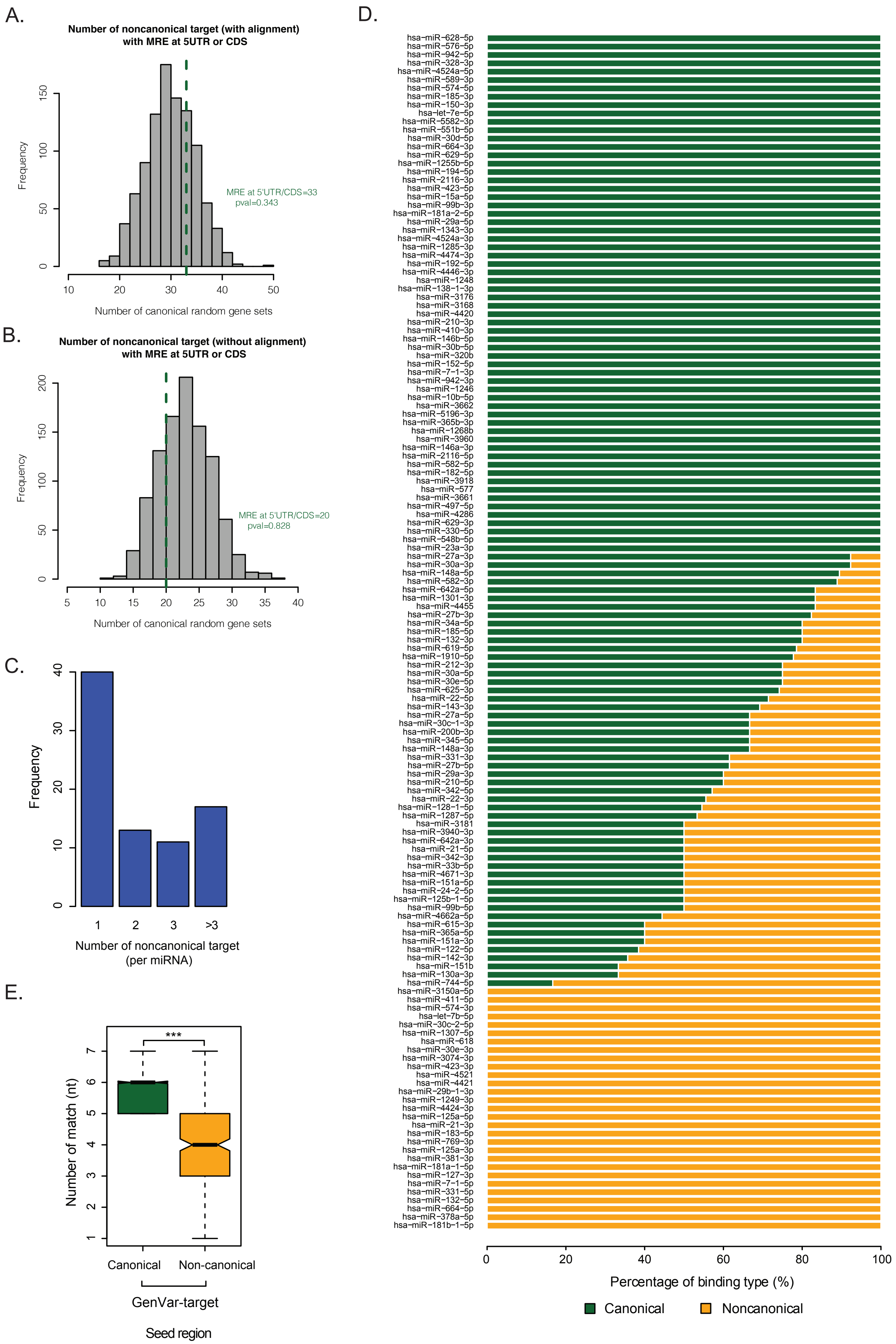
