## Supplementary Figure S3 for "Noncanonical targeting contributes significantly to miRNA-mediated regulation"

Supp Figure S3.

A.

| Prediction tools | # Targets | # Interactions | # miRNAs | # targets per miRNA | # miRNAs per target |
|---|---|---|---|---|---|
| GenVar-target | 593 | 665 | 143 | 5 | 1 |
| TargetScan | 13,842 | 474,114 | 143 | 3348 | 29 |
| RNA22 | 13,595 | 825,295 | 97 | 9209 | 60 |
| miRWalk | 13,027 | 448,787 | 111 | 4259 | 30 |
| All-expressed genes | 14,847 | | | | |

B.

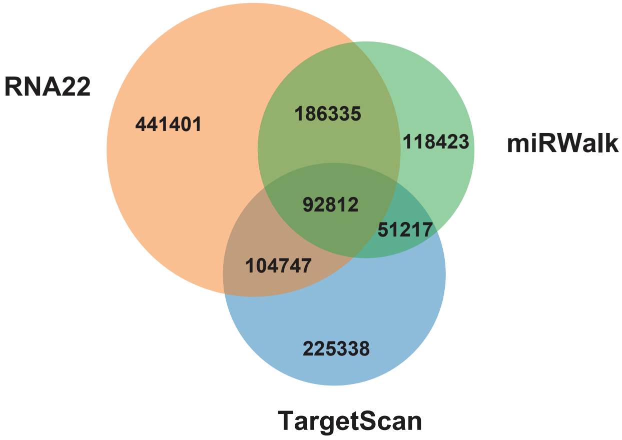
