## Supplementary Materials for "Noncanonical targeting contributes significantly to miRNA-mediated regulation"

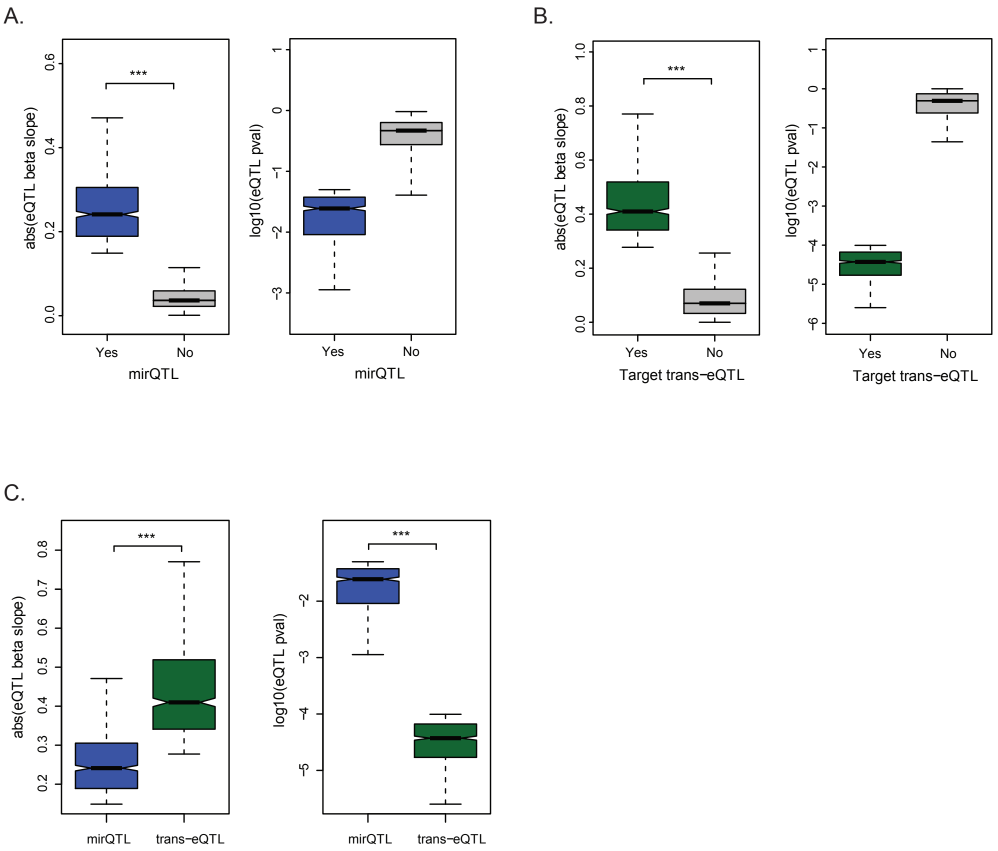


**Supplementary Figure 1. GenVar approach detects physiologically relevant miRNA targets.** Distribution of significant mirQTL (blue, A) and gene *trans*-eQTL (green, B) associations (right: beta slope; left: adjusted p-value) and those that are insignificant (grey) after multiple-testing correction. (C) Distribution of mirQTL (blue) and gene *trans*-eQTL (green) associations (right: beta slope; left: adjusted p-value). Differences between groups were tested using a two-tailed Mann-Whitney *U* test. *** $p < 0.001$.

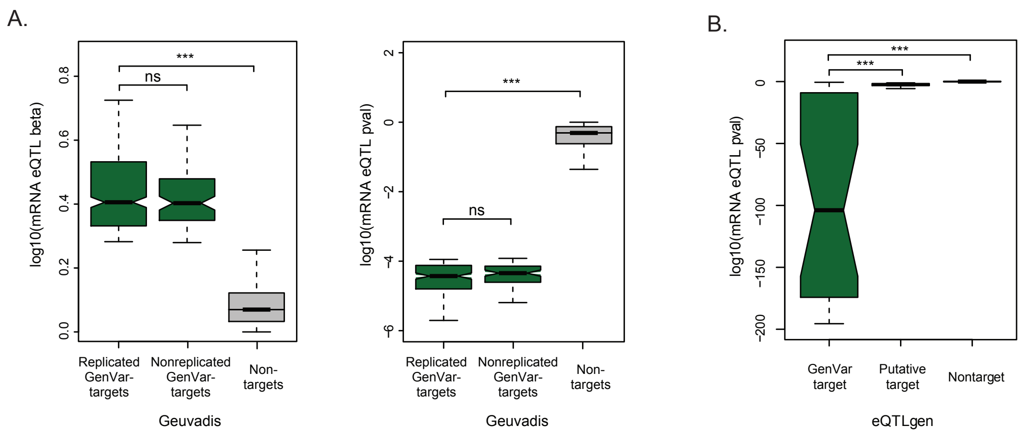


**Supplementary Figure 2. Limitations of the GenVar approach.** (A) Distribution of GenVar-target *tran*-eQTL associations with mirQTLs in Geuvadis LCLs (green, left: beta slope; right: adjusted p-value) that are replicated or non-replicated in eQTLgen blood samples, as well as mirQTLs associations with nontargets. (B) In eQTLgen blood samples, distribution of *trans*-eQTL associations (adjusted p-value) between validated GenVar-targets (dark green), putative targets (light green) or nontargets (grey) with mirQTLs. Differences between groups were tested using a two-tailed Mann-Whitney *U* test. * $p < 0.05$; ** $p < 0.01$; *** $p < 0.001$; NS $p > 0.05$.

A.

| Prediction tools | # Targets | # Interactions | # miRNAs | # targets per miRNA | # miRNAs per target |
|---|---|---|---|---|---|
| GenVar-target | 593 | 665 | 143 | 5 | 1 |
| TargetScan | 13,842 | 474,114 | 143 | 3348 | 29 |
| RNA22 | 13,595 | 825,295 | 97 | 9209 | 60 |
| miRWalk | 13,027 | 448,787 | 111 | 4259 | 30 |
| All-expressed genes | 14,847 | | | | |

B.

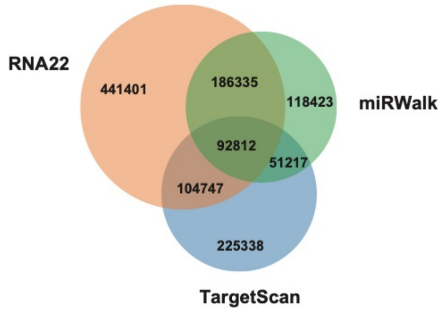


**Supplementary Figure S3. Comparison between GenVar-targets and predictions by other tools.** (A) Table of the number of miRNA target predictions by GenVar, TargetScan, RNA22, and miRWalk for LCL-expressed miRNAs and protein-coding genes. (B) Venn diagram of the overlap in predictions common to TargetScan, RNA22 and miRWalk.

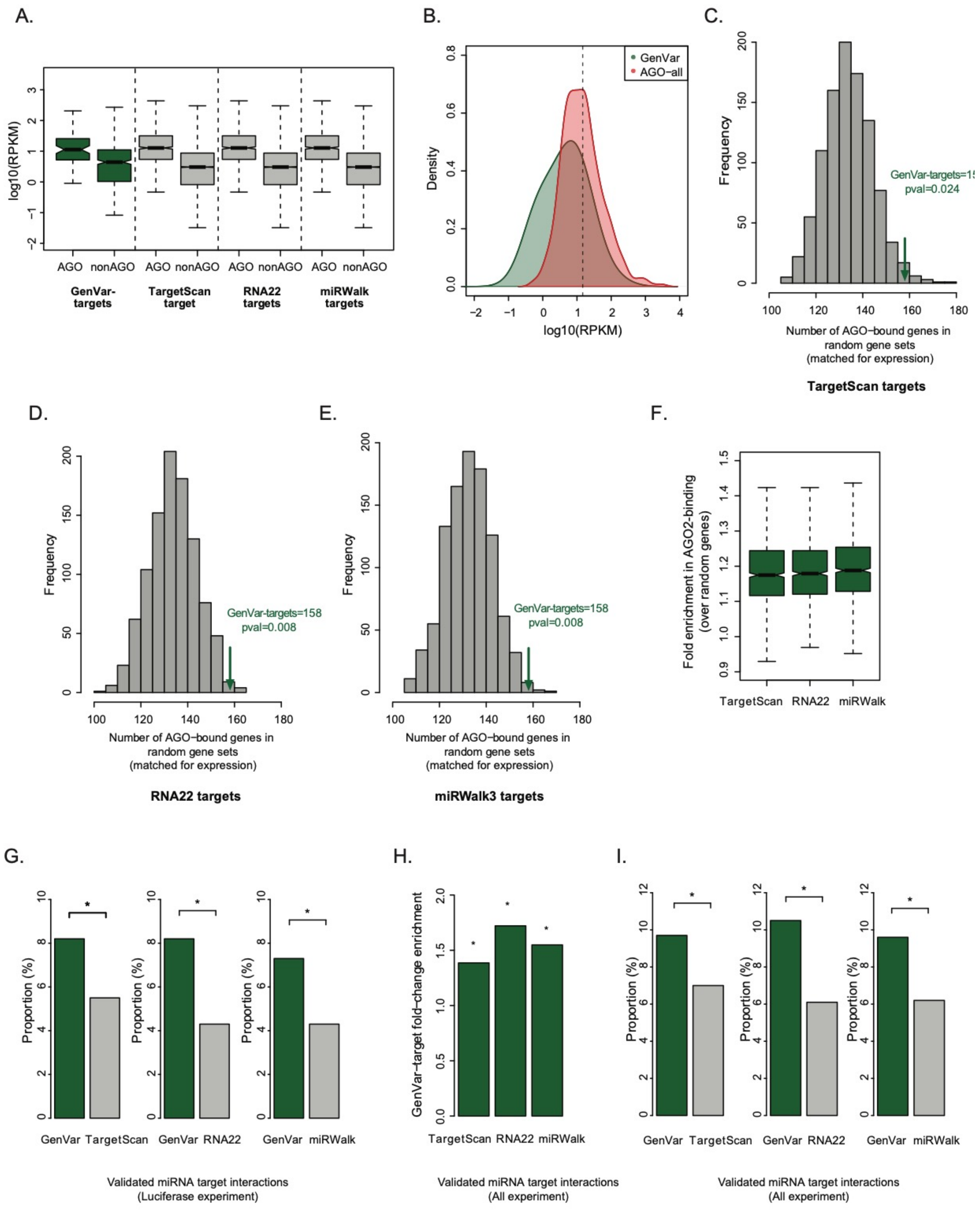


**Supplementary Figure 4. GenVar targets are a reliable set of miRNA targets.** (A) Distribution of expression levels (RPKM) of GenVar-target predictions (green) and predictions of genes by other tools (grey) of those bound and unbound by AGO2 in LCLs. (B) Histogram of the expression levels (RPKM) of GenVar-targets (green) and all genes bound by AGO2 (red). Distribution of AGO2-bound genes within 1000 sets of randomly sampled genes from (C) TargetScan, (D) RNA22, and (E) miRWalk with matching expression levels and 3' UTR length as GenVar-targets. Green arrow represents the number of AGO2-bound GenVar-targets. (F) Distribution of enrichment of the proportion of AGO2-bound GenVar-targets compared to 1000 sets of randomly sampled genes with matching expression predicted by

TargetScan, RNA22, or miRWalk. (G) Of all predictions by each of the three prediction tools, the proportion of experimentally validated (using direct luciferase assays) GenVar-target (green) predictions and nonGenVar-target predictions (grey). (H) Of all predictions by each of the three prediction tools, (H) the enrichment in the proportion and (I) the proportion of experimentally validated (using all catalogued experiments) GenVar-target predictions over nonGenVar-target predictions. Differences between groups were tested using a two-tailed Mann-Whitney *U* test (F,H) or two-tailed Fisher's exact test (G,I). * $p < 0.05$; ** $p < 0.01$; *** $p < 0.001$; NS $p > 0.05$.

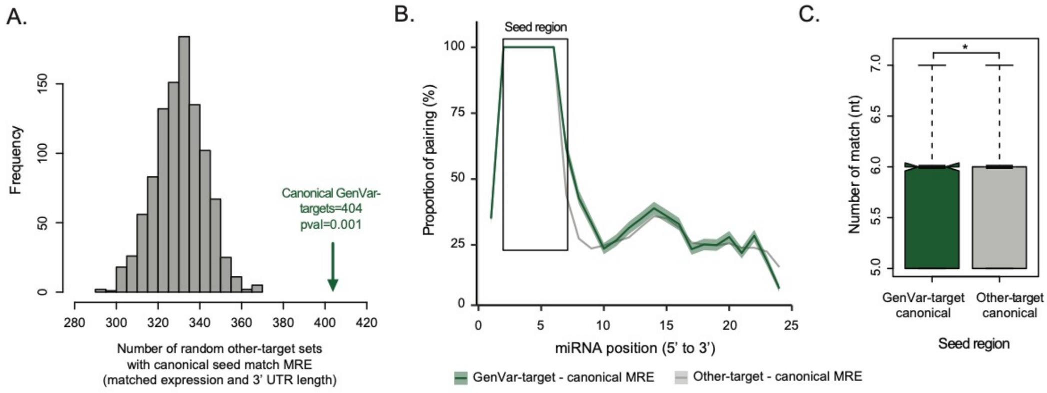


D.

| | Canonical binding interactions | | | |
|---|---|---|---|---|
| | Genvar-target | % | Other-target | % |
| 8mer | 109 | 17.3 | 192142 | 13.6 |
| 7mer | 190 | 30.2 | 345709 | 24.4 |
| 6mer | 331 | 52.5 | 879102 | 62.0 |

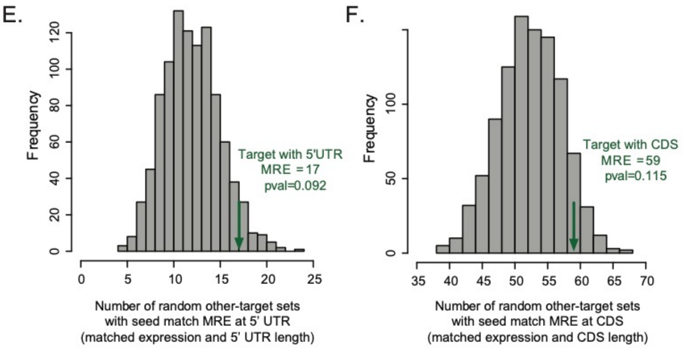


**Supplementary Figure 5. A sizeable fraction of GenVar-targets is noncanonical.** (A) Distribution of the number of genes with canonical MREs at their 3' UTRs within 1000 sets of randomly sampled genes with matching expression levels and 3' UTR length as GenVar-targets. Green arrow represents number of GenVar-targets with canonical MREs at their 3' UTRs. (B) Binding profile illustrating the average fraction of complementary sequence alignment between target and miRNA at each position across the body of the mature miRNA (5' to 3') for canonical GenVar-targets (green) and nontargets with canonical binding sites (grey). (C) Distribution of the number of aligned sequences (nucleotide) between GenVar-targets (green) or other predicted targets (grey) with the seed region of miRNAs. Differences between groups were tested using a two-tailed Mann-Whitney *U* test. * $p < 0.05$. (D) Table of the number and proportion of 8-mer, 7-mer, and 6-mer MREs within GenVar-target and other predicted target 3' UTRs. Distribution of the number of genes with canonical MREs at their (E) 5' UTRs and (F) CDS within 1000 sets of randomly sampled genes with matching expression levels and 5' UTR or CDS length as GenVar-targets, respectively. Green arrow represents number of GenVar-targets with canonical MREs at their 5' UTR or CDS.

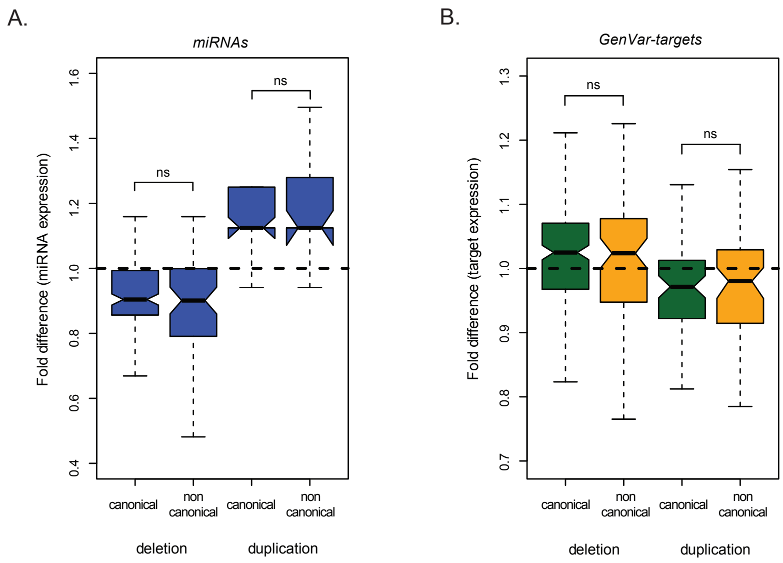


**Supplementary Figure 6. A sizeable fraction of GenVar-targets are noncanonical.** (A) Distribution of the fold difference in miRNA levels (CPM, blue), relative to the median of diploid samples, of individuals that carry homozygous deletions or multiple duplications at miRNAs that target canonical or noncanonical targets. (B) Distribution of fold difference in gene expression levels (RPKM) of canonical (green) or noncanonical (yellow) GenVar-targets, relative to the median of diploid samples, between individuals that carry homozygous deletions or multiple duplications at their targeting miRNAs. Differences between groups were tested using a two-tailed Mann-Whitney *U* test. NS $p > 0.05$.

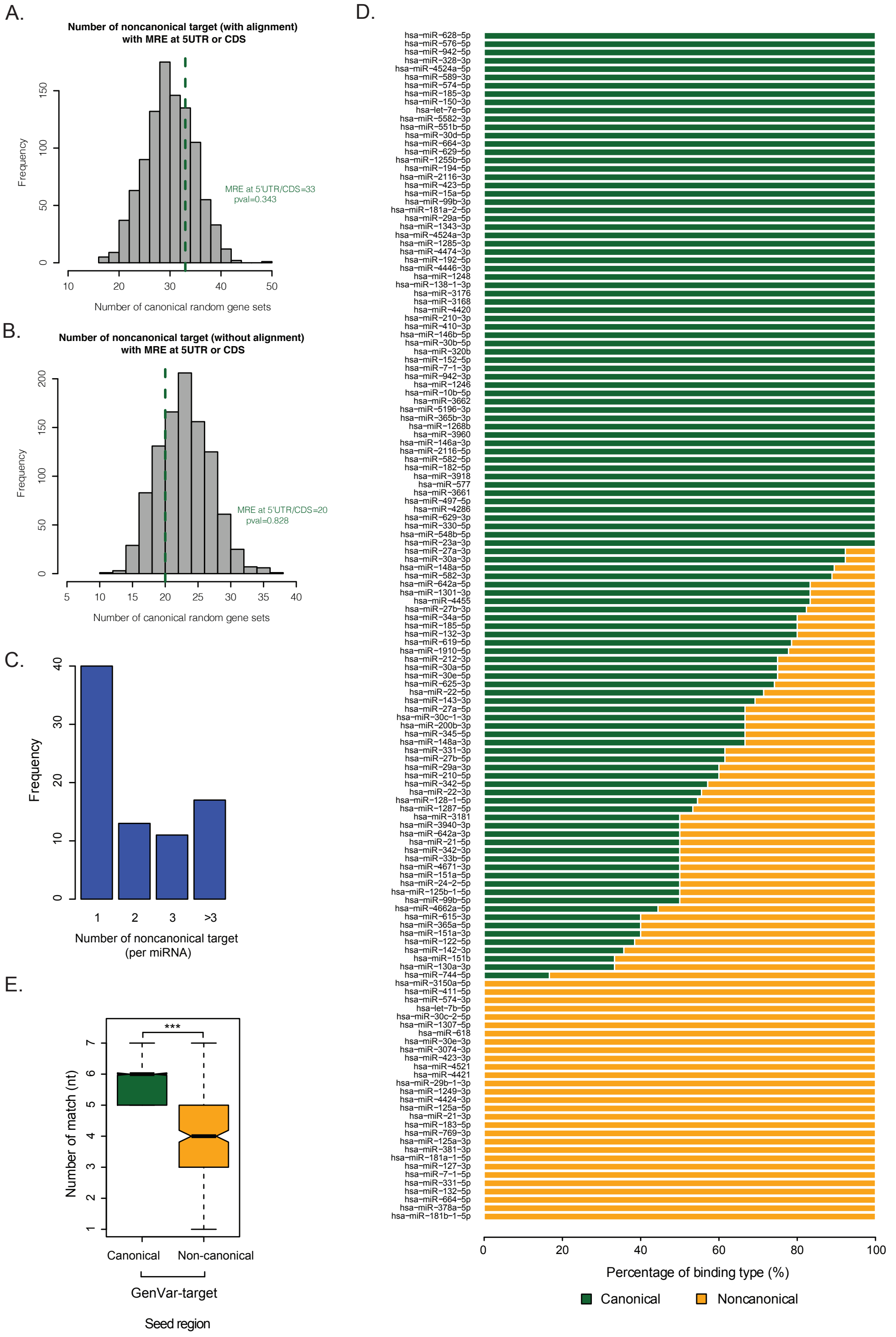


**Supplementary Figure 7. A sizeable fraction of GenVar-targets are noncanonical.** Distribution of the number of genes with canonical MREs at their 5' UTRs or CDS within 1000 sets of randomly sampled genes with matching expression levels and 5' UTR or CDS length as noncanonical GenVar-targets whose 3' UTRs (A) can or (B) cannot be aligned to their targeting miRNA. Green arrow represents number of noncanonical GenVar-targets with seed-matching MREs at 5'UTR or CDS for those whose 3' UTRs (A) can or (B) cannot be aligned to their targeting miRNAs. (C) Distribution of the number of noncanonical GenVar-targets per miRNA. (D) The proportion of canonical (green) and noncanonical (yellow) GenVar-targets for each miRNA. (E) Distribution of the number of complementary sequence alignments (nt) between canonical (green) or noncanonical (yellow) GenVar-targets with seed region of miRNAs. Differences between groups were tested using a two-tailed Mann-Whitney *U* test. *** $p < 0.001$.

### SUPPLEMENTARY TABLE LEGENDS

Supplementary Table ST1. Table of miRNA mirQTL and GenVar-target *trans*-eQTL associations.

Supplementary Table ST2. Copy number variations (CNVs) at primary miRNA transcript loci in TCGA blood-derived cancer samples (Lymphoid Neoplasm Diffuse Large B-cell Lymphoma, DLBC) with available genotyping and transcriptomic data.

Supplementary Table ST3. Table of canonical and noncanonical GenVar-targets per miRNA.
